# Vascular Deformation Mapping Calibration with Physics-based Synthetic Data on Multi-axial Aortic Motion

**DOI:** 10.64898/2026.05.20.726669

**Authors:** Taeouk Kim, Timothy Baker, Nicholas Burris, C. Alberto Figueroa

**Affiliations:** Department of Biomedical Engineering, University of Michigan, Ann Arbor, MI, USA; Department of Radiology, University of Michigan, Ann Arbor, MI, USA; Department of Radiology, University of Wisconsin–Madison, Madison, WI, USA; Department of Surgery, University of Michigan, Ann Arbor, MI, USA

**Keywords:** 3D aortic deformation, Aortic CTA, Image registration, rigidity penalty

## Abstract

Aortic stiffness is both heterogenous and anisotropic. Current non-invasive methods to estimate aortic stiffness are limited to characterizing the aortic tissue as isotropic due to the lack the techniques required to extract multi-axial strain from 3D dynamic images. Vascular deformation mapping (VDM) is a nonrigid image registration technique which has thus far been applied to map aortic growth using longitudinal imaging. In this study, we propose to use VDM to assess 3D aortic deformation by mapping diastolic and systolic images. During image registration process, penalty parameters are employed to fine-tune image alignment and penalize non-physiological deformations. These penalty parameters must be calibrated to ensure that VDM successfully reproduces multi-axial aortic motion patterns in health and disease. In this paper, we developed a calibration pipeline for these parameters using synthetic data. A rotation-free shell model was used to generate physics-based synthetic data on aortic motion incorporating patient-specific geometries, root motion, and blood pressure from a cohort of 14 subjects (healthy, Marfan’s syndrome and thoracic aortic aneurysm). An error metric was defined to quantify the quality of the VDM results. Furthermore, a k-means clustering technique was used to categorize the subjects into three clusters based on ascending aortic motion. Optimal penalty parameters were identified for each of the three clusters. The results indicated that patient clusters with smaller aortic root motion required larger rigidity penalty values. The calibrated parameters successively reduced errors in 3D displacement and multi-axial stretch compared to un-optimized VDM predictions, enhancing the accuracy of capturing aortic deformation from dynamic images. Among the different aortic regions, the ascending thoracic aorta exhibits the largest error reduction.

## 1. Introduction

Aortic stiffness is both heterogeneous, displaying spatially-varying values in different regions along the aorta, and anisotropic, indicating distinct stiffness patterns in axial and circumferential directions [1, 2, 3, 4]. Aortic stiffness has been recognized as an important indicator of vascular health [5, 6]. Typically, older patients or those with aortic diseases, such as aortic dissection or Marfan’s syndrome, exhibit stiffer aortas [7, 8, 9, 10, 11, 12, 13]. Numerous efforts have been made to estimate aortic stiffness. Some stiffness calibration efforts have relied on invasive approaches, whereby strips of aortic tissue from different regions are tested in both axial and circumferential directions [14, 15, 16, 17]. However, invasive methods have been limited because they require surgical aortic tissue samples and can only be conducted in specific regions. As an alternative, non-invasive, dynamic image-based approaches have been used to quantify aortic stiffness and strains. However, most of these approaches have been limited to isotropic descriptions of the tissue and have thus been unable to discriminate stiffness and strains in different spatial directions [18, 19, 20]. These non-invasive approaches have either relied on 1D or 2D descriptions of aortic motion. Therefore, there is a pressing need to develop methods for assessing 3D aortic deformation from dynamic 3D images, to estimate multi-axial aortic stiffness and strain.

Vascular deformation mapping (VDM) is a nonrigid image registration technique, originally developed to assess 3D aortic growth by using longitudinal computed tomography angiography (CTA) data acquired at two different times during clinical surveillance (herein, we will refer to this technique as VDM(G): VDM “growth”) [21, 22]. When performing nonrigid image registration on dynamic ECG-gated CTA (i.e. registration between images within the cardiac cycle), it is possible to obtain a description of 3D deformation. We will refer to this technique as VDM(D) : VDM “deformation “. Both VDM(G) and VDM(D) use the Elastix open-source software to perform nonrigid image registration [23]. In Elastix, penalty parameters are employed to fine-tune image alignment and penalize non-physiological deformations. These penalty parameters include bending energy, affine, orthonormality and properness [24]. Figure 1 shows a VDM(D) workflow where registration is performed between two images within the cardiac cycle, corresponding to diastole and systole. The output is a 3D displacement field over the entire aorta which makes it possible to calculate 3D strains. The penalty parameters must be calibrated to ensure that VDM(D) successfully reproduces multi-axial aortic motion patterns in health and disease.

**Figure 1.**
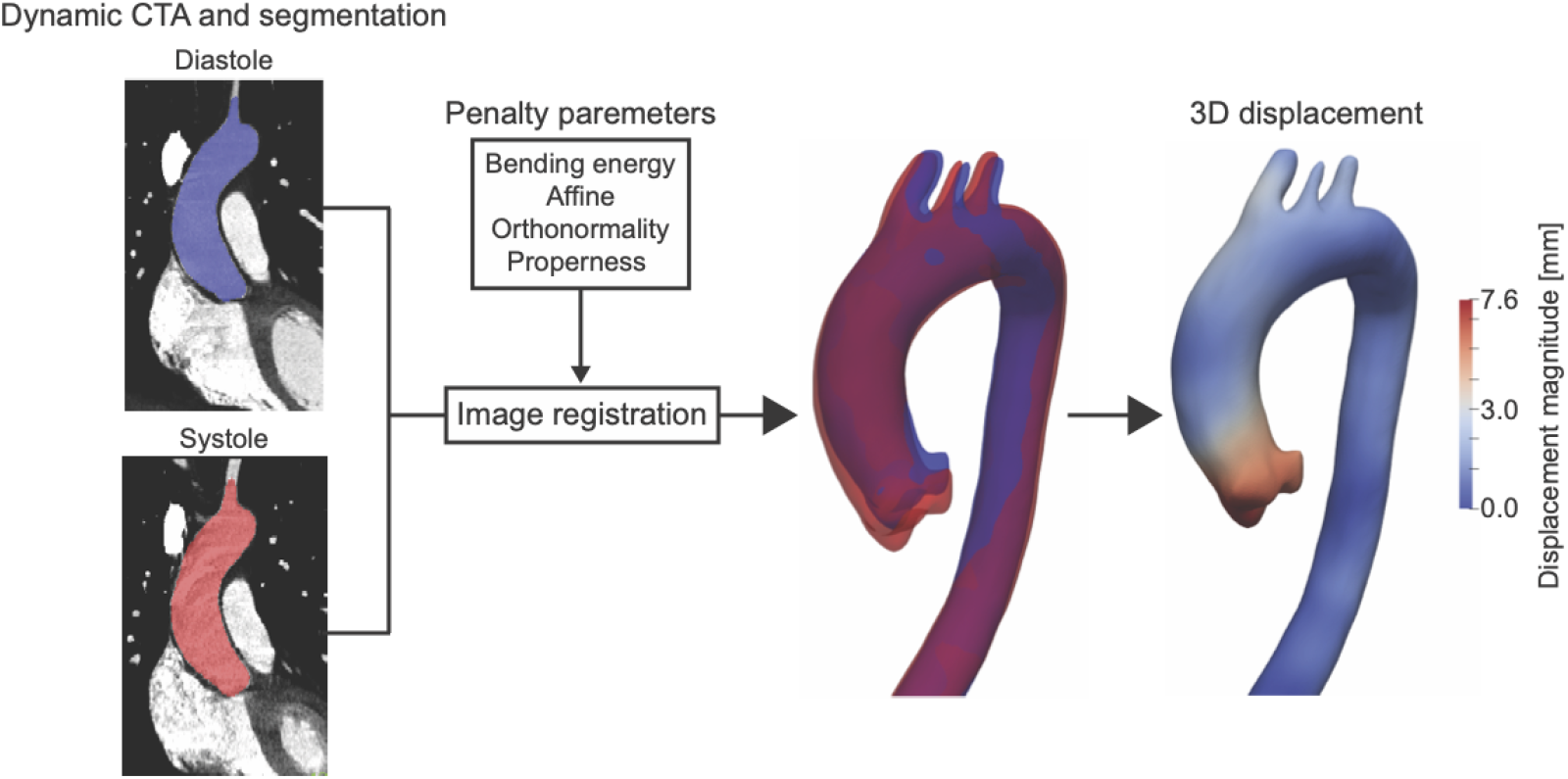
VDM(D) workflow.

These penalty parameters serve to regularize and constrain the registration but lack a direct physics-based definition; therefore, their tuning to ensure accurate and physiologically realistic motion patterns is imperative. For example, non-calibrated penalty parameters may lead to non-physical 3D displacement fields. This problem is illustrated in Figure 2. The top row shows synthetic data on 3D aortic displacement, as well as axial and circumferential stretches generated using a non-linear shell model and realistic pressure and root displacement boundary conditions (details described in the methods section). Here, the axial direction is tangent to the center-line, while the circumferential direction is perpendicular to the centerline. The bottom row shows results of VDM(D) performed with non-calibrated penalty parameters, taking as inputs the systolic and diastolic configurations of the synthetic data. Even though the reconstructed 3D displacement looks qualitatively similar to the ground truth data, the stretches (particularly the axial stretch) reveal profound differences. This behavior is not unexpected, because stretches are derivatives of the displacement and thus errors in the displacement are amplified accordingly. This example highlights the importance of calibrating VDM(D) penalty parameters to ensure physically meaningful deformation maps.

**Figure 2.**
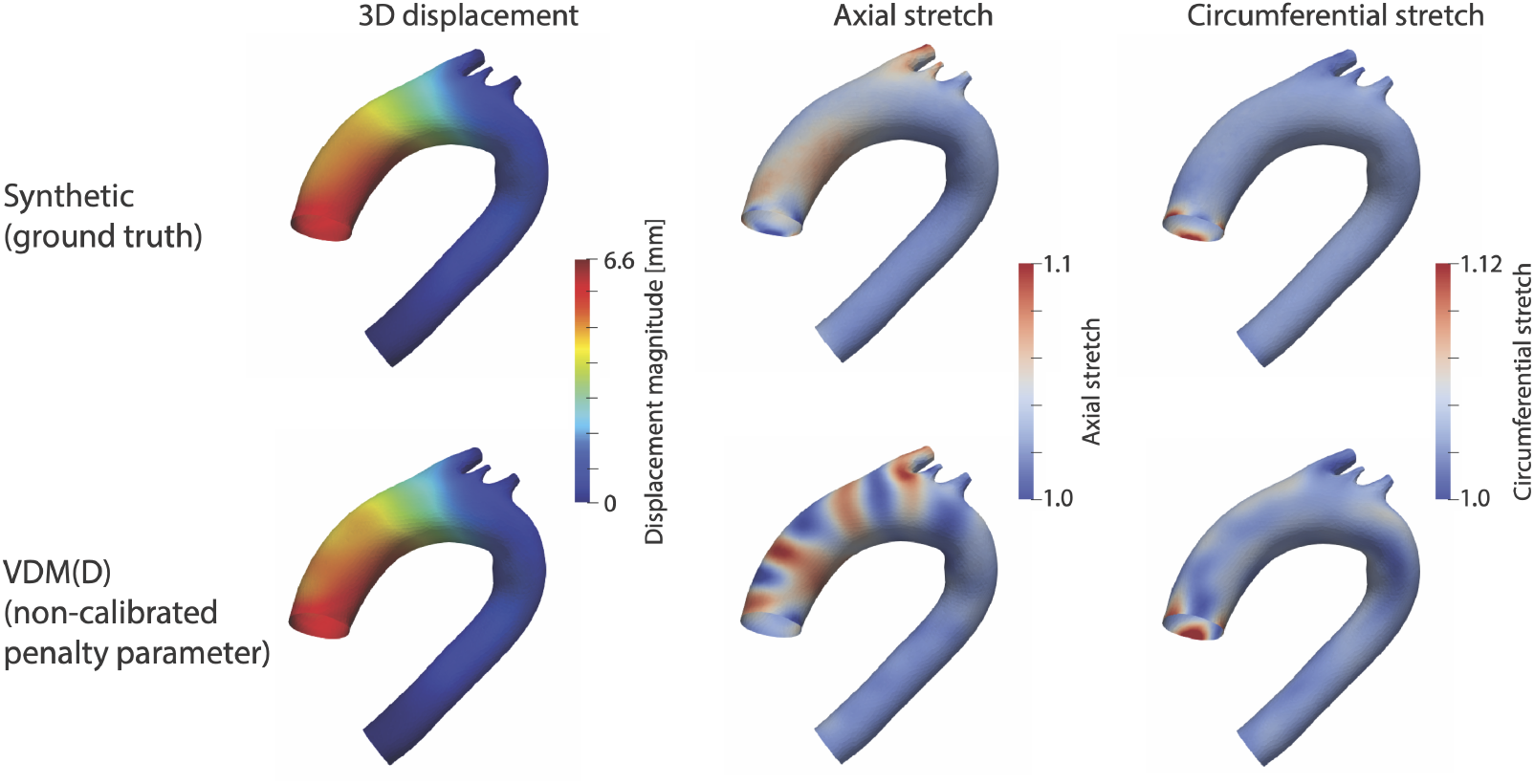
3D displacement and multi-axial stretch analysis using synthetic data (top) and reconstructed using VDM(D) with non-calibrated penalty parameters (bottom).

A key challenge in calibrating VDM(D) penalty parameters is the absence of established ground truths for 3D displacement across the entire aorta. Most prior art has been limited to either 1D descriptions of aortic root motion [25, 26] or 3D data at only specific locations down the aorta [27, 28]. Therefore, aortic displacement from these methods is not sufficient to validate and calibrate the four-dimensional (i.e., volumetric and time-resolved) nonrigid registration performed in VDM(D). To address this issue, in this paper we generated synthetic data on aortic motion for the purpose of calibrating VDM(D), using physiologically realistic data on pressure and root motion for a total of 14 different aortas representing healthy and diseased conditions.

## 2. Methods

### 2.1. Penalty parameters

The Elastix (and thus VDM(D)) registration algorithm maximizes the voxel-wise mutual information between a fixed diastolic image and a deformed systolic image. To constrain the nonrigid transformation to be anatomically plausible, we incorporated two regularization terms: a rigidity penalty *P*_rigid_ that enforces locally rigid behavior within specific image regions, and a bending-energy penalty *P*_BE_ that ensures smoothness of the deformation field.

Formally, VDM(D) minimizes the cost function

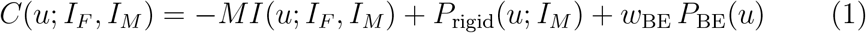

where *MI* is the mutual information between the fixed image *I*_*F*_ (*x*) : Ω ⊂ ℝ^*d*^ →ℝ and the moving image *I*_*M*_ (*x*) : Ω ⊂ℝ^*d*^→ ℝ after applying the displacement field *u*(*x*) using the nonrigid transformation *T* (*x*) = *x* + *u*(*x*). The other terms in (1) are penalty terms that constrain the optimization.

VDM(D) adopts the rigidity penalty *P*_rigid_ introduced by Staring *et al*. (2007) [24] and implemented in `Elastix`. Following their formulation, a transformation is locally rigid if it is (i) affine in *x*, (ii) has an orthonormal local Jacobian, and (iii) has a determinant of +1. These three conditions respectively define the affine constraint (AC), orthonormality constraint (OC), and properness constraint (PC). For clarity, we summarize the 2D versions of these terms here, whereas VDM(D) utilizes their 3D counterparts.

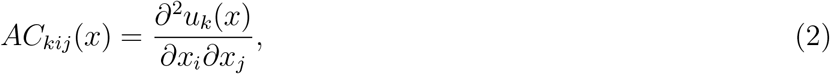

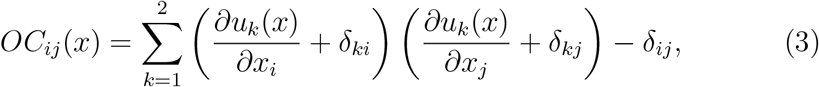

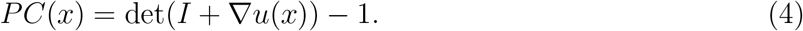

for all ∈{*i, j, k* 1, 2}, not counting duplicates. Deviations from these ideal conditions are penalized according to the rigidity penalty

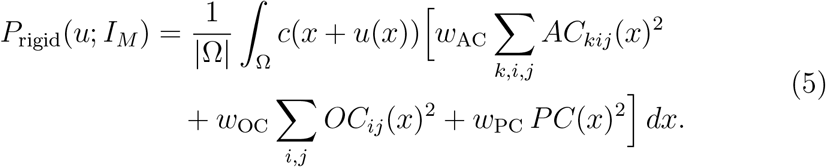

where *c*(*x*) ∈ [0, 1] defines which image regions are rigid (*c* = 1) or deformable (*c* = 0), and *w*_AC_, *w*_OC_, and *w*_PC_ are weights that determine the relative importance of each rigidity subterm. VDM(D) sets *c*(*x*) = 1 for voxels located within the aortic segmentation mask and *c*(*x*) = 0 for all other voxels.

To further regularize the deformation field, VDM(D) employs the bending energy (BE) penalty introduced by Rueckert *et al*. (1999) [29] and also implemented in `Elastix`. BE penalizes the second-order spatial derivatives of the deformation field, discouraging abrupt curvature

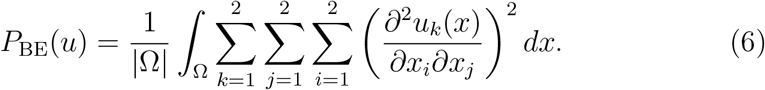

Overall, BE encourages smooth deformations, whereas the AC, OC, and PC constraints enforce local rigidity, yielding more realistic registration results [24].

Each penalty term has an associated weight that controls its influence on the optimization. The ranges of weights tested are based on prior research [24] and our experiments: the BE weight *w*_BE_ was varied from 0 to 220, the AC weight *w*_AC_ from 0 to 550, the OC weight *w*_OC_ from 0 to 1.5, and the PC weight *w*_PC_ from 0 to 1.5. Total 144 combinations of weights were tested using a four-dimensional grid search scheme (see Figure 3). Figure 3 shows one example of error values (defined in section D. Error calculation) across different combinations of penalty parameters. The layout is organized as a 3×4 grid of subplots, where each subplot corresponds to a unique pair of the BE weight (*w*_BE_; columns) and the AC weight (*w*_AC_; rows). Within each subplot, a 3 ×4 table reports the error of the PC weight (*w*_PC_; rows) and OC weight (*w*_OC_; columns).

**Figure 3.**
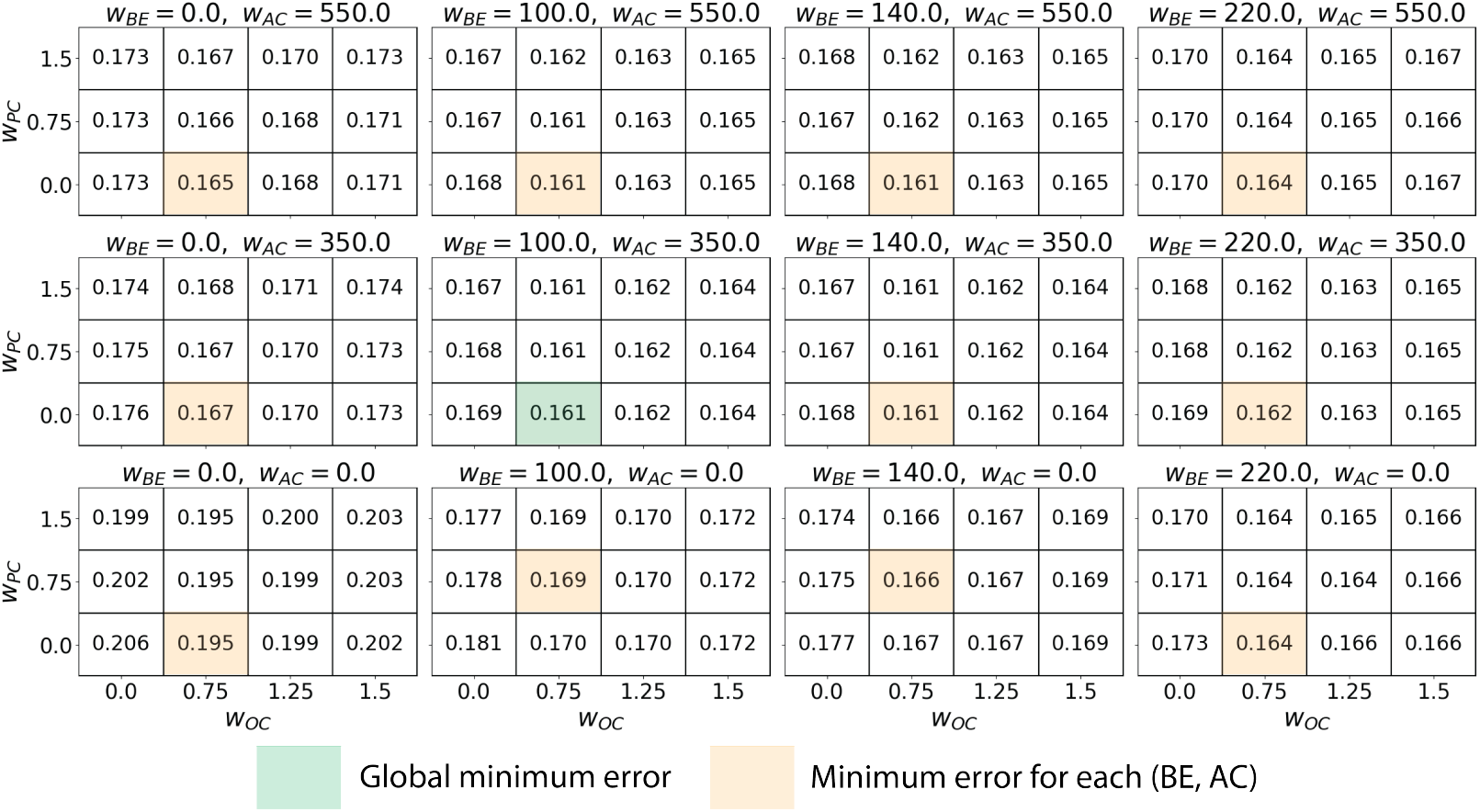
Error values with different penalty parameter combinations. Green box shows global minimum error and yellow box shows local minimum error for each BC and AC pair.

### 2.2. Synthetic data generation

A key challenge in calibrating VDM(D) penalty parameters is the absence of established ground truths for three-dimensional displacement across the entire aorta. Direct measurement of aortic tissue mechanical properties invivo remains limited. Ex-vivo mechanical testing can provide detailed material characterization, but it requires surgical tissue samples and is therefore restricted to a limited patient population, with additional methodological variability. Non-invasive imaging-based approaches, such as ultrasound or tagged MRI, can estimate regional deformation but are not widely available for the entire aorta and are subject to measurement noise and reconstruction errors.

Because of these limitations, we employed a physics-based simulation framework to generate synthetic data with known ground-truth displacement fields. This approach enables systematic calibration and evaluation of the VDM(D) penalty parameters under controlled conditions. We used a non-linear rotation-free shell formulation with population-based heterogeneous aortic stiffness.

#### 2.2.1. Rotation-free shell

A non-linear rotation-free shell formulation, solved using the finite element method to produce 3D displacement over the aortic surface, was used to generate the synthetic data [30]. This formulation is validated and has been used to handle large deformations for applications in vascular biomechanics. An incompressible Neo-Hookean constitutive model was considered for the aortic wall, as it is suitable for describing the nonlinear stress-strain behavior under large deformations. Stiffness parameters used for this model are described in the following section 2.2. The density and thickness of the aorta were set to 1000 kg/m^3^ and 2.0 mm, respectively [31, 32]. Boundary conditions for the aortic model are presented in Figure 4 (a). For the inlet face, we prescribed patient-specific 3D aortic root motion [7], while the arch and outlet faces were fixed. Patient-specific blood pressure was applied inside of the aortic wall.

**Figure 4.**
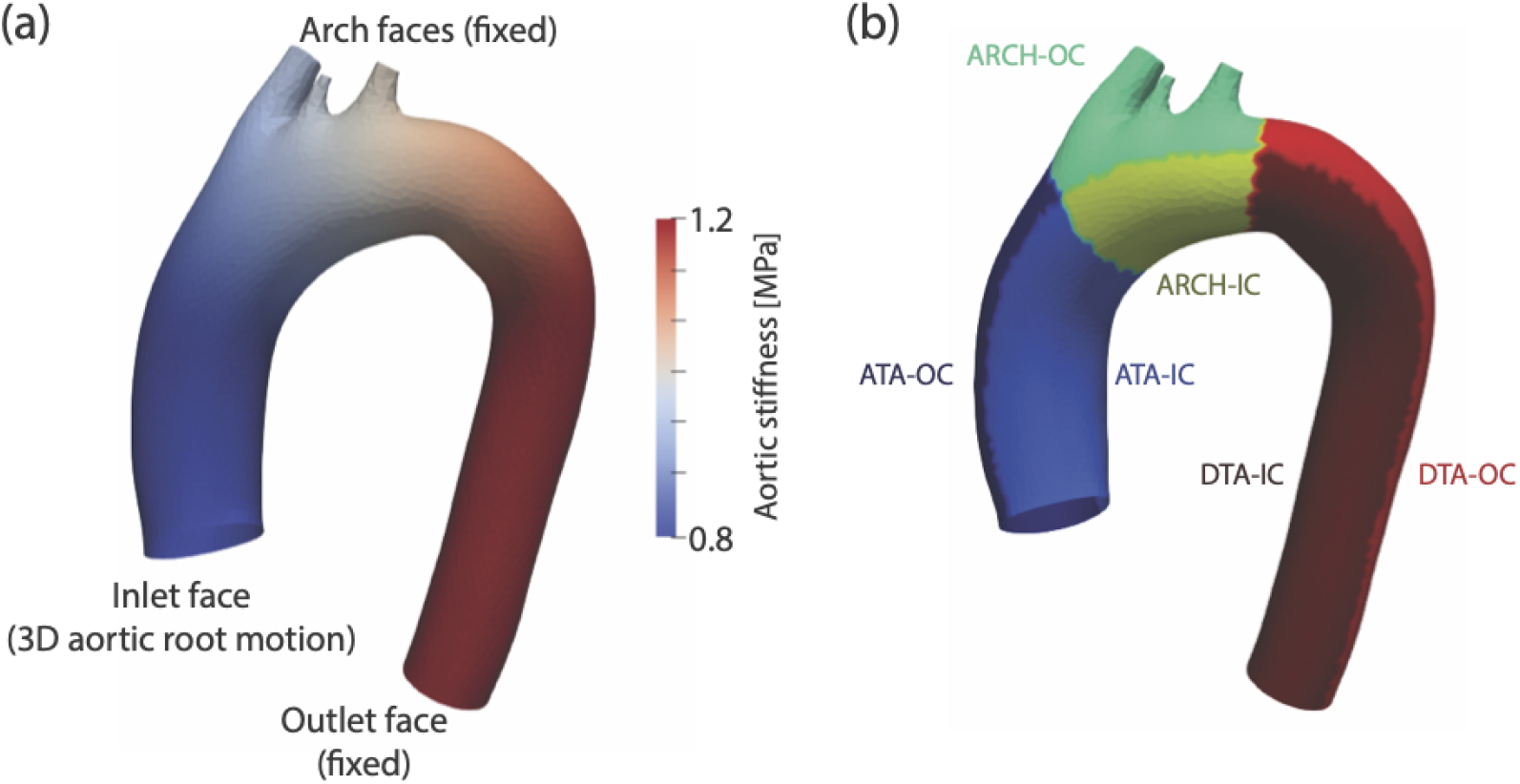
(a) Boundary conditions for synthetic data generation and heterogeneous aortic stiffness. (b) Six subregions for the error calculation. ATA: ascending thoracic aorta, DTA: descending thoracic aorta, IC: inner curvature, and OC: outer curvature.

#### 2.2.2. Population-based heterogeneous aortic stiffness

The proper choice of aortic stiffness is important for dynamic aortic simulations. In this paper, we used population-based heterogeneous aortic stiffness values. Our cohort includes non-aneurysmal patients as well as those with degenerative thoracic aortic aneurysms (TAA) and Marfan syndrome (Marfan). To determine baseline stiffness, we first extracted age-based stiffness data for the ascending thoracic aorta (ATA) and descending thoracicaorta (DTA) from the literature, which reports both axial and circumferential stiffnesses for non-aneurysmal patients [33]. Since we employed an isotropic model, we averaged the axial and circumferential stiffness values. To adjust for TAA and Marfan patients, we applied stiffness ratios of 1.26 and 1.65 on the ATA stiffness, respectively, based on differences from non-aneurysmal patients [7]. ATA and DTA stiffness values were linearly interpolated on the aortic surface. Figure 4 (a) shows one example of the heterogeneous aortic stiffness values on the aortic surface. ATA and DTA stiffness values for each patient can be found in Table 1.

**Table 1.** Patient characteristics and aortic stiffness. BP: blood pressure; NA: non-aneurysmal.

| Disease Category | Age | BP [mmHg] | ATA stiffness [MPa] | DTA stiffness [MPa] | 3D Root motion [mm] | Cluster |
| --- | --- | --- | --- | --- | --- | --- |
| NA 1 | 65 | 107/61 | 0.834 | 1.152 | 7.1 | 1 |
| NA 2 | 63 | 164/98 | 0.834 | 1.152 | 6.6 | 1 |
| NA 3 | 32 | 100/62 | 0.386 | 0.737 | 8.6 | 1 |
| TAA 1 | 66 | 139/78 | 1.05 | 1.152 | 6.9 | 1 |
| TAA 2 | 43 | 116/73 | 0.486 | 0.737 | 3.0 | 0 |
| TAA 3 | 75 | 142/77 | 1.05 | 1.152 | 3.8 | 0 |
| TAA 4 | 67 | 124/85 | 1.05 | 1.152 | 4.8 | 0 |
| TAA 5 | 64 | 110/72 | 1.05 | 1.152 | 6.5 | 1 |
| TAA 6 | 69 | 131/85 | 1.05 | 1.152 | 7.0 | 1 |
| Marfan 1 | 36 | 133/66 | 0.637 | 0.737 | 10.0 | 2 |
| Marfan 2 | 64 | 114/66 | 1.376 | 1.152 | 3.0 | 0 |
| Marfan 3 | 52 | 115/68 | 0.637 | 0.737 | 8.4 | 1 |
| Marfan 4 | 42 | 96/54 | 0.637 | 0.737 | 11.1 | 2 |
| Marfan 5 | 40 | 125/85 | 0.637 | 0.737 | 7.6 | 1 |

### 2.3. Synthetic image generation

A synthetic systolic image and mask were generated from the displacement field computed by rotation-free shell. An Elastic Body Spline (EBS) transformation was applied to the diastolic image using the vertices of the diastolic aortic surface and the synthetic systolic aortic surface as two sets of landmark points. EBS transforms the diastolic image into a synthetic systolic image by mapping corresponding landmark points to one another while interpolating intermediate points in a manner consistent with a physical model of a homogeneous, isotropic 3D elastic body [34]. EBS transformation is applied using Elastix’s Transformix program [23] with a Poisson ratio of 0.3.

### 2.4. Error calculation

To quantitatively evaluate the performance of VDM(D) with different penalty parameters, we first calculate a 3D displacement error field by comparing the VDM(D) results with the ground truth from the rotation-free shell model. Next, we define an appropriate error metric from the 3D displacement error field. Our error metric calculation involves three steps. 1) We define six subregions of aorta to ensure equal contributions from all aortic surfaces. The aorta is first segmented into three regions ATA, arch, and DTA - based on side vessel configuration. Each of these regions is then further divided into two subregions: inner curvature (IC) and outer curvature (OC). Figure 4 (b) illustrates these six subregions, including ATA-IC, ATA-OC, ARCH-IC, ARCH-OC, DTA-IC, and DTA-OC. 2) For each subregion, we calculate the maximum displacement error, and the root mean square error to evaluate local and global errors, respectively. A subregion error is average of these two errors. 3) The error for aorta is calculated by averaging the six subregion errors. In summary, error metric is defined as

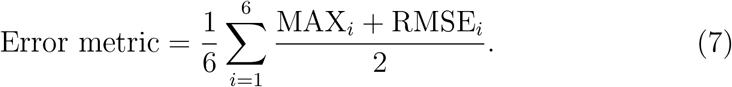

where *i* indexes the regions, MAX_*i*_ is the maximum displacement error in region *i*, and RMSE_*i*_ is the root-mean-square error.

### 2.5. VDM(D) penalty parameter calibration pipeline

Figure 5 shows our VDM(D) penalty parameter calibration pipeline. (1) Synthetic data generation: We calculated a 3D displacement of aorta from diastole to systole using the rotation-free shell formulation with an incompressible Neo-Hookean constitutive model. This defines the ground truth for our pipeline. (2) Synthetic image generation: we generated a synthetic systolic image and mask. The computed displacement field on the aortic surface from step (1) was interpolated through the 3D aortic volume to generate a synthetic systolic medical image and corresponding mask using EBS transformation. (3) VDM(D) analysis: the VDM(D) with penalty parameters was run to calculate 3D displacement from the reference diastolic and the synthetic systolic images and their masks. (4) Error calculation: VDM(D) displacements are compared against the computational ground truth, and the discrepancy was quantitatively calculated by using the error metric defined in the previous section ‘Error calculation method’. (5) Calibration of penalty parameters: we change the penalty parameters based on the error. We repeated steps (3), (4), and (5) with different penalty parameters until we found a minimal error.

**Figure 5.**
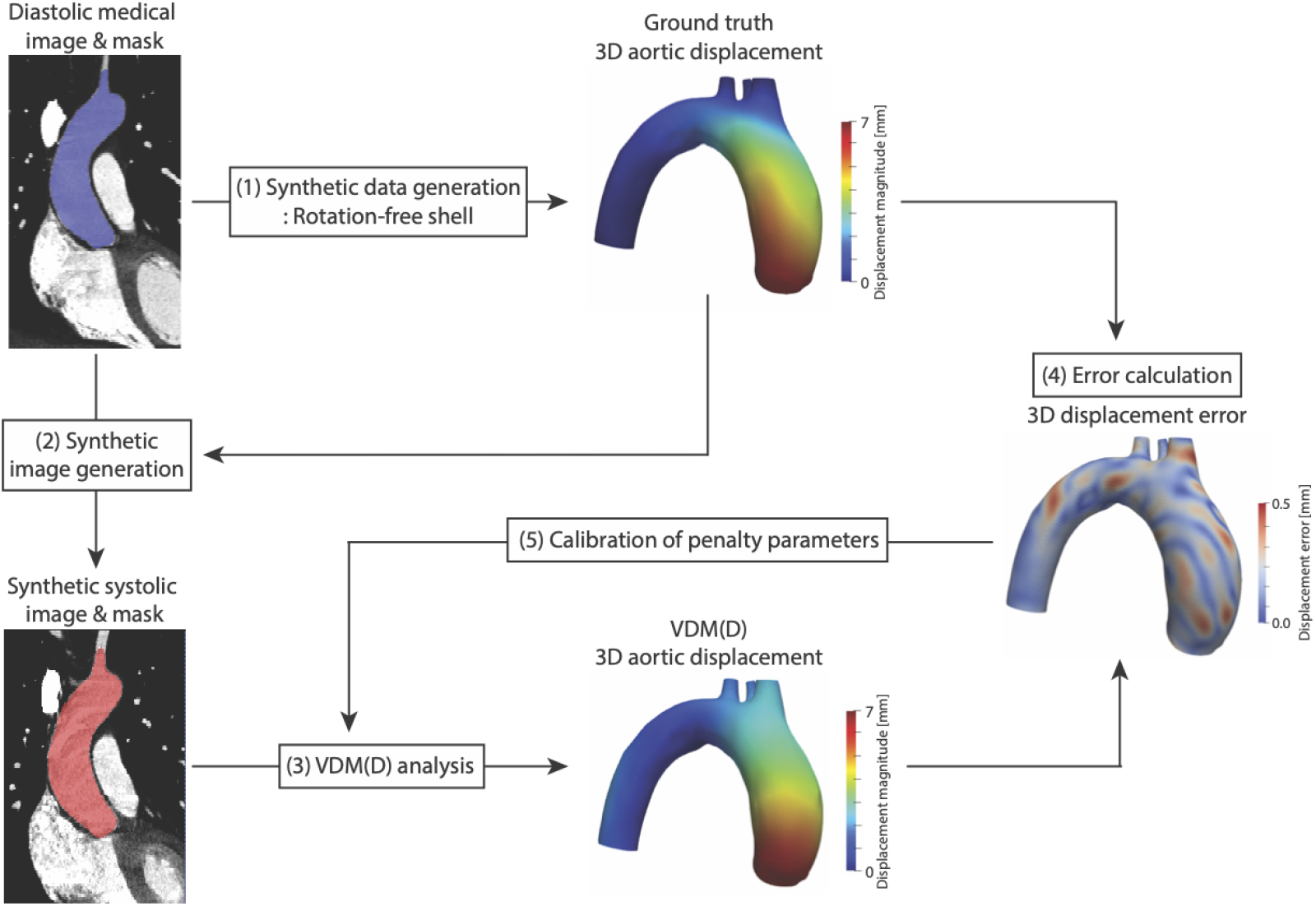
VDM(D) penalty parameters calibration pipeline.

### 2.6. Patient data

All procedures were approved by the local institutional review board, and a waiver of informed consent was obtained for this retrospective study. The study was conducted in compliance with the Health Insurance Portability and Accountability Act (HIPAA). Fourteen patients, categorized into three groups (non-aneurysmal, TAA, and Marfan), were selected based on the availability of high-quality, multi-phase ECG-gated CTA data. The peak systolic and end-diastolic phases of the ECG-gated CTA imaging were selected and extracted using open-source software, OsiriX MD [35]. Dynamic CT imaging covered the entire thoracic aorta (i.e., proximal arch vessels through the diaphragm). Details of the patient data can be found in Table 1. There were 3, 6, and 5 patients from the non-aneurysmal, TAA, and Marfan groups, respectively. Overall, the mean age was 55.4 ±14.4 years. The average systolic and diastolic blood pressures were 122.6 ±18.3 mmHg and 73.6 ±11.9 mmHg, respectively. ATA and DTA stiffness values were calculated following the previous section “Population-based heterogeneous aortic stiffness”. The 3D root motion magnitude was measured as described in our previous publication [7].

### 2.7. Subject classification strategies

VDM(D) penalty parameters were calibrated using three distinct grouping methods, leading to the identification of global, disease-specific, and motion-specific penalty parameters. The global penalty parameter was identified as the one that minimized error across all fourteen patients. In contrast, disease-specific and motion-specific penalty parameters were tailored for each disease group (non-aneurysmal, TAA, and Marfan) and for each cluster divided by root motion, respectively. We employed the k-means clustering technique for classification based on root motion to avoid arbitrary thresholds when separating clusters [36]. K-means clustering is an algorithm that groups data points into K number of clusters, where each data point belongs to the cluster with the nearest average. In this study, we used 3 clusters. The cluster 0, 1, and 2 include patients with root motion less than 5 mm, between 5 and 9 mm, and larger than 9 mm, respectively. Each cluster includes 4, 8, and 2 patients. The clustering results can be found in Table 1. The penalty parameters for each group were determined by minimizing the error across the patients within that group.

## 3. Results

### 3.1. Penalty parameters

Table 2 shows the global, disease-specific, and motion-specific penalty parameters obtained from our calibration pipeline. Overall, the ranges of calibrated penalty parameters are between 100-220, 0-350, 0.75-1.5, and 0.75-1.5 for the BE, AC, OC, and PC, respectively. While no specific trends are observed across different disease groups, there is a correlation between motion-specific penalty parameters and aortic root motion. Patient clusters with smaller aortic root motion require higher penalty values. For example, cluster 0 shows largest penalty parameter values: BE/AC/OC/PC = 220/350/1.5/1.5, while cluster 2 shows the smallest penalty parameter values: BE/AC/OC/PC = 100/0/0.75/0.75.

**Table 2.** Global, disease-specific, and motion-specific penalty parameters. NA: non-aneurysmal.

| Category | Groups | BE | AC | OC | PC |
| --- | --- | --- | --- | --- | --- |
| Global | – | 140 | 0 | 0.75 | 1.5 |
| Disease-specific | NA | 100 | 350 | 0.75 | 0.75 |
|  | TAA | 140 | 0 | 1.5 | 1.5 |
|  | Marfan | 100 | 0 | 0.75 | 0.75 |
| Motion-specific | Cluster 0 | 220 | 350 | 1.5 | 1.5 |
|  | Cluster 1 | 140 | 0 | 0.75 | 1.5 |
|  | Cluster 2 | 100 | 0 | 0.75 | 0.75 |

### 3.2. Error metric values with different penalty parameters

Figure 6 shows the error metric values without penalty parameters (represented by blackyellow bar) and with three different penalty parameter sets (represented by blackblue, blackred, and blackmagenta bars for global, disease-specific, and motion specific, respectively). Nearly all patients show that penalty parameters help reduce error compared to no penalty. The average errors across all patients are 0.211, 0.167, 0.164, and 0.163 for the no penalty, global, disease-specific, and motion specific, respectively. Given that the motion-specific penalty parameters are obtained from the objective grouping method and result in the lowest error, our final choice among the grouping methods is the motion-specific penalty parameters. More details about the parameter choice will be described in Discussion 2. Penalty parameter choice. From here, results using penalty parameters implies the use of motion-specific penalty parameters. The largest reduction in error metric achieved with penalty parameters, compared to no penalty, is 44.3% for patient 6. The average error reduction is 20.4%.

**Figure 6.**
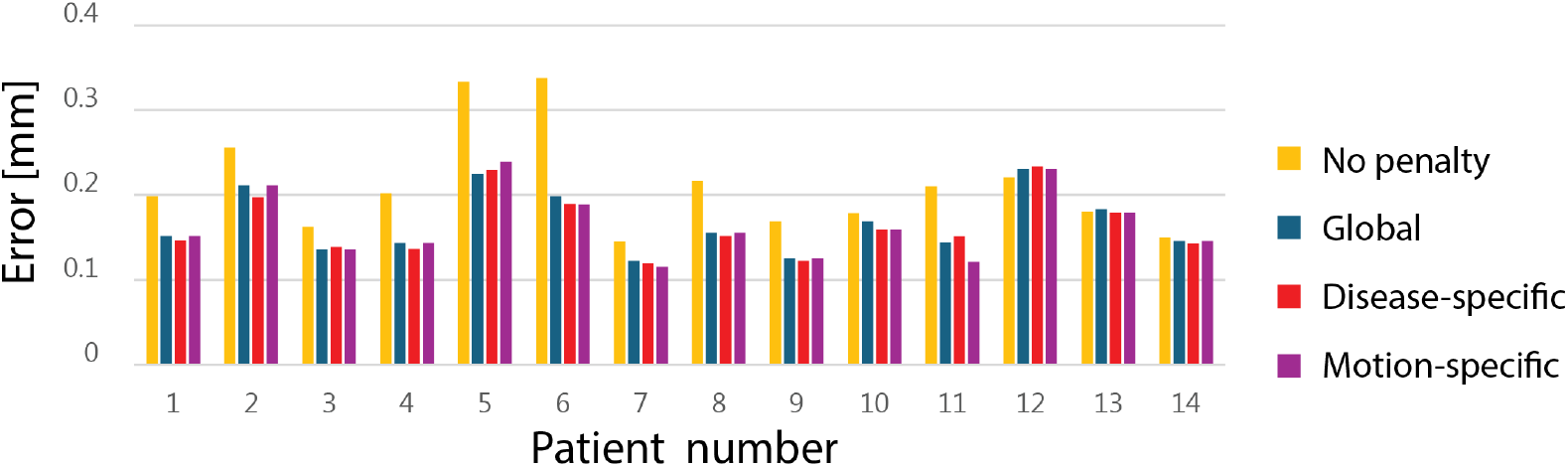
Error metric for each patient with different penalty parameters: no penalty, global, disease-specific, and motion-specific.

### 3.3. Surface error plots with and without penalty parameters

Figure 7 illustrates the surface error plots, which shows the 3D displacement error between ground truth from the rotation-free shell and VDM(D) results, both without and with penalty parameters applied. Four aortas are presented: two representing the best cases (patient 5 and 6 from Figure 6) and two representing the worst cases (patient 12 and 13 from Figure 6). Two aortas from the best error improvement show that properly calibrated penalty parameters successively reduce high displacement errors. The largest error decreases from 0.74 mm to 0.31 mm for the first patient and 0.58 mm to 0.26 mm for the second patient. Physiologically spurious bands in error maps are also smoothed out by using penalty parameters (see second aorta of the best error improvement). Two aortas from the worst error improvement show relatively small displacement error without penalty (maximum error of 0.24 mm and 0.38 mm) compared to the best error improvement aortas without penalty (maximum error of 0.74 mm and 0.58 mm). These aortas show no improvement or a slight increase in error with penalty application.

**Figure 7.**
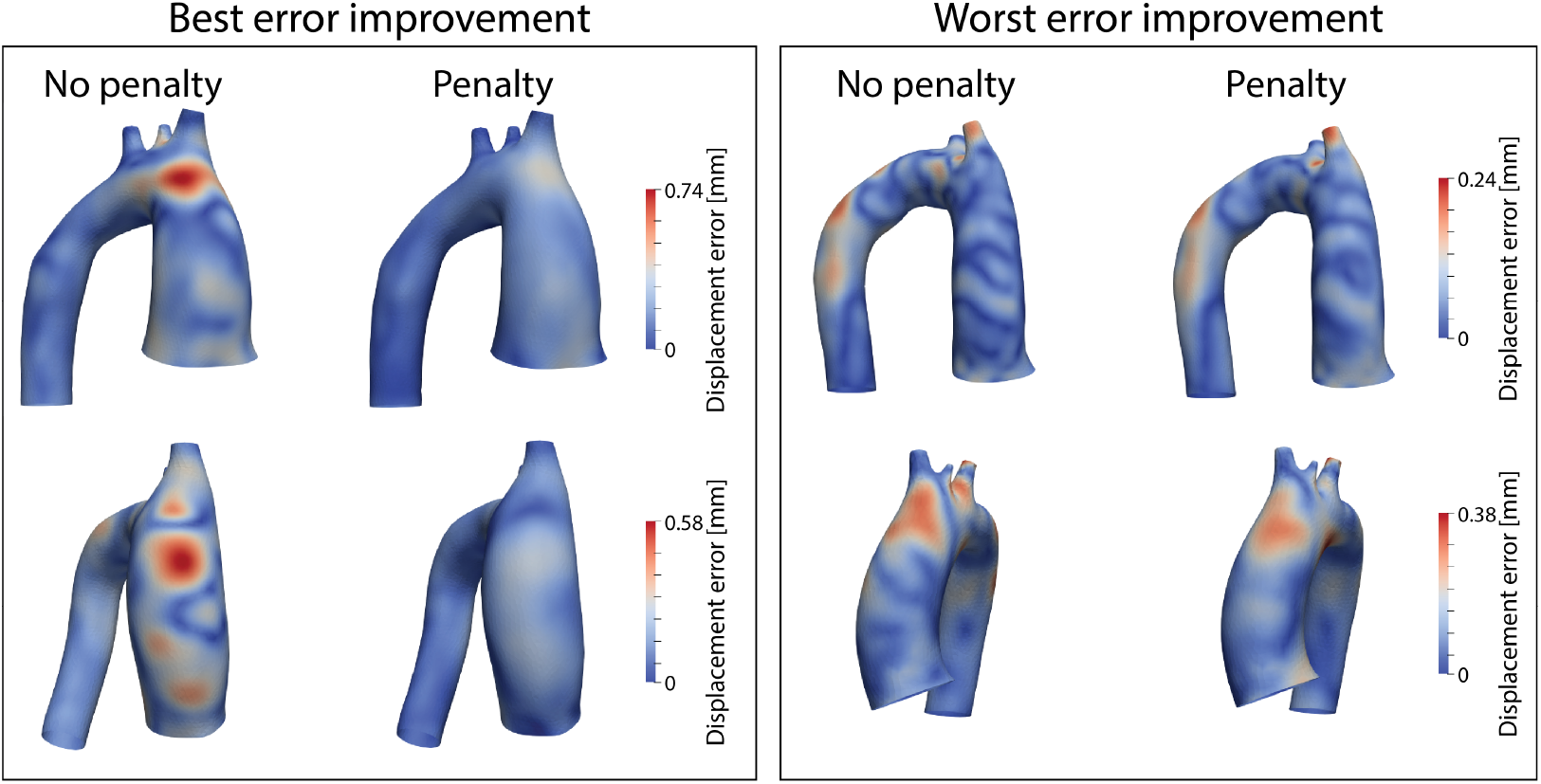
3D displacement error analysis: comparison between scenarios with no penalty and with calibrated penalty, including the two most improved and two least improved error cases.

## 4. Discussion

### 4.1. Stretch comparison

Figure 8 compares 3D displacement and multi-axial stretch between the ground truth (derived from the rotation-free shell) and VDM(D) results using different penalty parameter values. The axial direction is tangent to the centerline, while the circumferential direction is perpendicular to the centerline. The comparison between ground truth and VDM(D) with non-calibrated penalty parameters emphasizes that even small errors in 3D displacement can result in large errors in multi-axial stretch, particularly in the axial direction (as indicated by the red star in Figure 8). Among the multi-axial aortic stretches, axial stretch derives primarily from downward pulling of the ascending aortic as a result of left ventricular contraction, whereas circumferential stretch results from inflation due to blood pressure. The synthetic 3D displacement field in Figure 8 indicates that overall ascending aortic deformation is primarily influenced by relatively large and complex root motion, more so than inflation. This complex motion challenges VDM(D)’s ability to accurately capture deformation without properly calibrated penalty parameters, resulting in a higher error in axial stretch compared to circumferential stretch.

**Figure 8.**
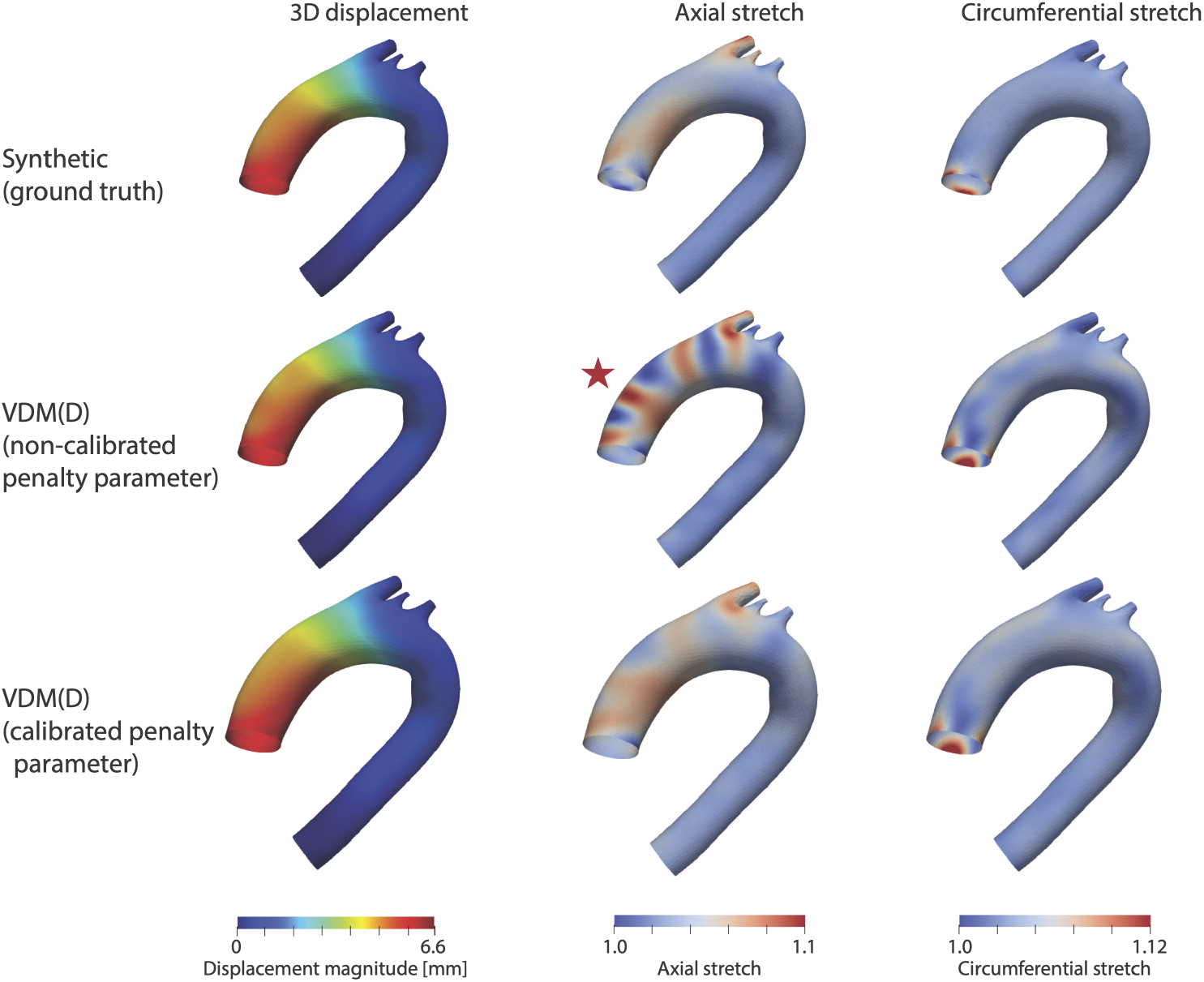
3D displacement and multi-axial stretch comparison between ground truth (top) and VDM(D) results with different penalty parameter values: non-calibrated (middle) and calibrated using our pipeline (bottom).

The 3D displacement magnitudes at the red star in the Figure 8 are 4.62 mm for the ground truth and 4.64 mm for VDM(D) with non-calibrated parameters. This minor difference of 0.02 mm results in a significant discrepancy in axial stretch values, which are 1.021 and 1.079, respectively. This sensitivity arises from the definition of stretch, which is a derivative of the 3D displacement in specific directions. With properly calibrated penalty parameters, the VDM(D) results show a significant reduction in stretch discrepancy compared to the ground truth, particularly in the axial direction. The axial stretch at the red star region is 1.033 with calibrated penalty parameters. These results demonstrate that properly calibrated VDM(D) effectively captures deformation from the root motion. The multi-axial strain obtained from non-invasive medical images can be used to estimate the multi-axial aortic stiffness.

### 4.2. Penalty parameter choice: disease-specific vs motion-specific

In this study, we used two different grouping methods: disease-specific and motion-specific for optimizing parameter selection. The disease-specific method categorized patients into non-aneurysmal, TAA, and Marfan groups. Non-aneurysmal and TAA patients were classified based on a maximum diameter threshold of 40 mm, without any syndromic features and no family history of aortic diseases. The aortic diameter was collected from clinical radiology reports. The Marfan syndrome group clarification was established using the Ghent criteria, which incorporate aortic root dilatation and FBN1 mutations in the diagnosis [37, 38]. However, the challenges of using maximum diameter or genetic characteristics for parameterizing the VDM(D) analysis is that these diameter thresholds can be somewhat arbitrary and are variable due to measurement imprecision, and further the genetic diagnosis of an individual is often not known with confidence, and thus such a disease etiology based parameterization is undesirable due to its subjective nature [39, 40, 41, 42, 43].

In contrast, root motion provides a more objective metric and more directly reflects the magnitude of aortic deformation. The magnitude of deformation is a key factor influencing deformable image registration performance, as the registration algorithm is agnostic to the underlying disease state and instead tracks voxel intensity changes between images. Therefore, grouping patients based on root motion better aligns the penalty parameter selection with the deformation characteristics encountered by the registration algorithm. However, the primary challenge with using root motion for patient classification is selecting the appropriate threshold between different groups. The k-mean clustering technique offers an objective threshold based on the given data. Since disease-specific and motion-specific penalty parameters showed similar performance (see Figure 6), we chose to use motion-specific method due to its more objective grouping approach. The similar performance of disease-specific and motion-specific penalty parameters may be attributed to the fact that the disease groups inherently exhibit different root motions in our cohorts. The average root motions for the motion-specific penalty parameters may be attributed to the inherent differences in root motion among the disease groups in our cohort. The average root motions for the non-aneurysmal, TAA, and Marfan groups were 7.4 (±1.0), 5.3 (±1.0), and 8.0 (±3.1) mm, respectively (p-value: 0.063, ANOVA). These differences in root motion may already influence the values of disease-specific penalty parameters.

### 4.3. Correlation between penalty parameter and root motion

Motion-specific penalty parameters demonstrate a correlation with the magnitude of root motion: clusters with smaller root motion need larger penalty parameter values (see Table 2). When root motion is smaller, the deformation of the aorta between diastole and systole behaves in a more rigid fashion. The penalty parameters employed in this study guide deformable image registration, resulting in an overall increase in registration rigidity. A larger rigidity penalty imposes a stronger constraint on deformation, favoring more rigid alignments, which is appropriate for the small root motion. In contrast, a lower rigidity penalty allows for larger deformation, enabling the registration algorithm to accommodate larger magnitude and/or more complex motions, including non-rigid deformations. This is useful when aligning images that undergo large changes in shape or when there is significant motion like axial motion due to cardiac contraction [44, 45]. This flexibility from using different penalty parameter values is particularly important for aortic image registration, as the thoracic aorta experiences a wide range of deformations influenced by root motion, varying with age and disease [7].

### 4.4. Correlation between error and root motion

The error also correlates with root motion. For in-depth analysis, we split the 3D root motion into axial and in-plane directions. The axial motion is defined as the projection of the 3D root motion in the direction of the tangential of the centerline. Conversely, the in-plane motion is defined as the projection of the 3D root motion in the direction perpendicular to the centerline (see Figure 9). Different motion values are provided in Table 3. The average root motions were 6.79 (±2.48), 5.41 (±2.13), and 4.41 (±1.78) mm for the 3D, axial, and in-plane directions, respectively.

**Table 3.** 3D, axial, and in-plane root motions for all patients.

| Patient | 3D root motion [mm] | Axial root motion [mm] | In-plane root motion [mm] |
| --- | --- | --- | --- |
| NA 1 | 7.1 | 6.32 | 3.21 |
| NA 2 | 6.6 | 5.81 | 4.72 |
| NA 3 | 8.6 | 5.51 | 6.95 |
| TAA 1 | 6.9 | 4.97 | 5.20 |
| TAA 2 | 3.0 | 2.57 | 1.82 |
| TAA 3 | 3.8 | 2.02 | 3.68 |
| TAA 4 | 4.8 | 4.35 | 2.13 |
| TAA 5 | 6.5 | 5.09 | 4.57 |
| TAA 6 | 7.0 | 5.59 | 4.28 |
| Marfan 1 | 10.0 | 8.06 | 7.81 |
| Marfan 2 | 3.0 | 2.28 | 2.26 |
| Marfan 3 | 8.4 | 7.38 | 4.51 |
| Marfan 4 | 11.1 | 9.27 | 6.23 |
| Marfan 5 | 7.6 | 6.56 | 4.44 |

**Figure 9.**
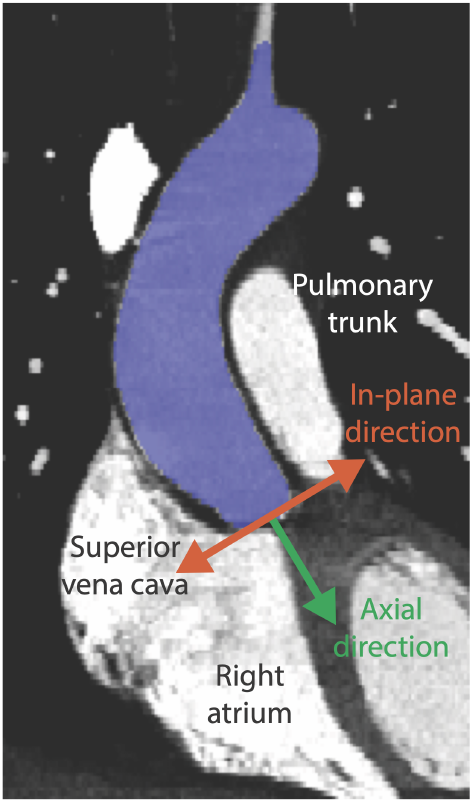
Axial and in-plane directions of the root motion with anatomies near aorta.

Figure 10 illustrates the nine different correlations between error and various root motions. Correlations between variables were assessed using linear regression. A p-value *<* 0.05 was considered indicative of a strong correlation. The ‘kruskalwallis’ and ‘fitlm’ functions in MATLAB were used for the Kruskal–Wallis test and linear regression, respectively. Plots (a), (b), and (c) show the correlations between errors from VDM(D) without using penalty parameters and three types of root motion. There is a strong negative correlation with 3D root motion and axial motion, with p-values of 0.034 and 0.031, respectively, while no significant correlation is observed with in-plane motion. These results suggest that VDM(D) without penalty parameters can effectively register two aortas with large deformations, while it does not have enough rigidity to accurately capture small deformations. These correlations are particularly strong in the ATA region (see Figure 4 (b)). Plots (d), (e), and (f) depict the correlation between errors, which were calculated only for the ATA region using VDM(D) without penalty parameters, and three types of root motions. These plots demonstrate smaller p-values and larger magnitudes of R values than those in plots (a), (b), and (c). This indicates that the effect of root motion is stronger in the ATA than in the arch and DTA regions. In-plane root motion shows weaker correlations with error (p-values *>* 0.05) compared to 3D and axial root motions. This can be attributed to the relatively smaller magnitude on in-plane motion and its easier capture by VDM(D), as the aorta is surrounded by other structures such as the superior vena cava, right atrium, and pulmonary trunk in the in-plane direction (see Figure 9).

**Figure 10.**
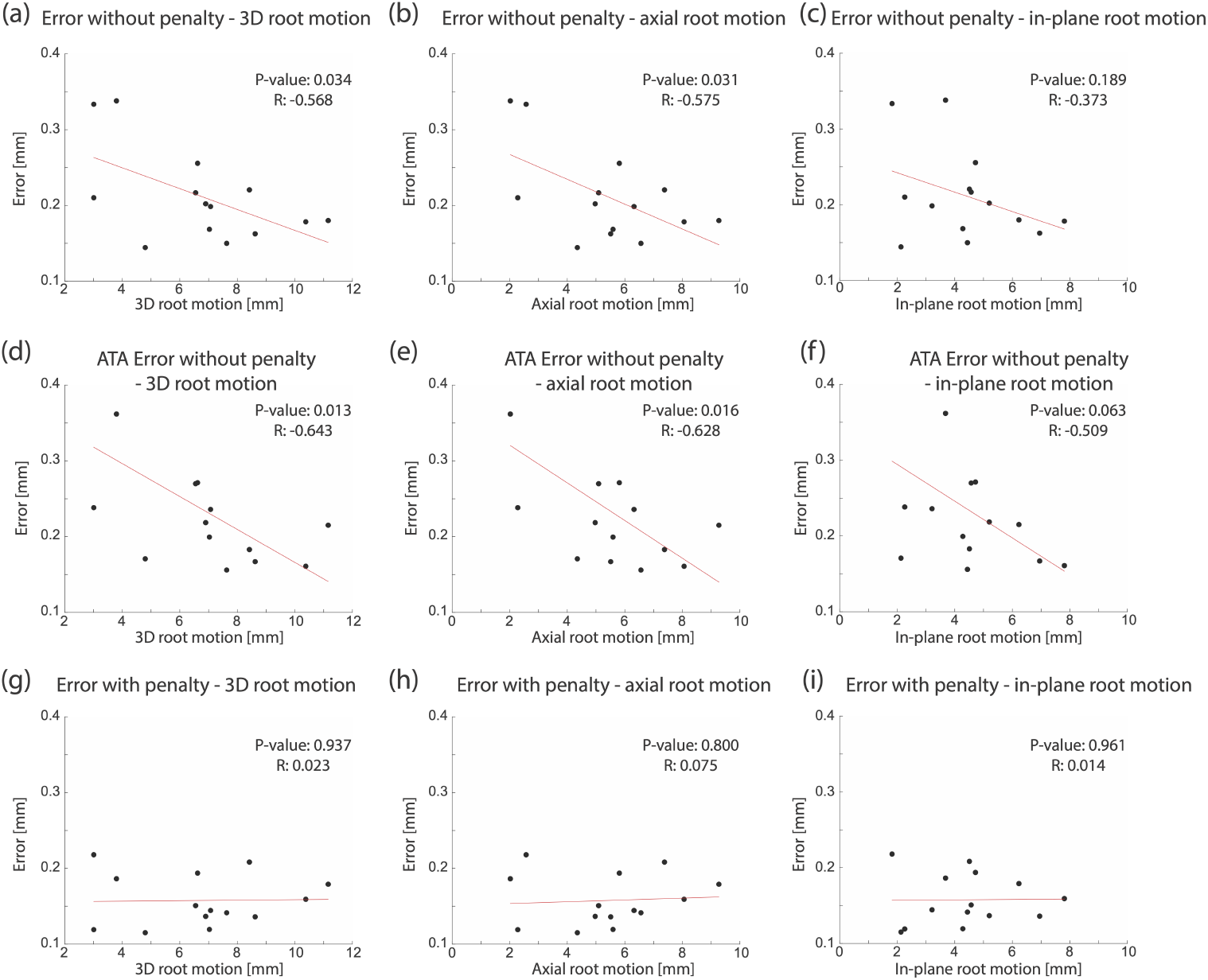
Correlations between errors and three types of root motion. (a), (b), (c): Error without penalty. (d), (e), (f): ATA Error without penalty. (g), (h), (i): Error with penalty. (Red line shows the line of the regression.

### 4.3. Homogenous penalty parameter

In this study, we implemented the homogenous penalty parameters. However, different aortic regions experience varied deformations. Figure 11 (a) shows the 3D displacement field on the aortic surface, where the ATA, arch, and DTA regions exhibit displacement ranges of 4–7 mm, 2–4 mm, and 0–2 mm, respectively. This variation in displacement distribution originates from cardiac motion that affects the root and ascending segments most significantly. In this study, we showed the correlation between optimal penalty parameters and aortic motion. Since we used a homogeneous penalty parameter on the aorta with heterogeneous motion, different regions exhibited different levels of error reduction. Figure 11 (b) demonstrates the 3D displacement error between the ground truth and VDM(D), both with and without penalty parameters. With penalty applied, the maximum displacement error in the ATA region decreased from 0.346 mm to 0.181 mm (47% decrease), whereas the arch and DTA regions decreased from 0.320 mm to 0.282 mm (11.8% decrease) and from 0.225 mm to 0.200 mm (11.1% decrease), respectively. These results indicate that a penalty parameter tuned for large motion is effective in the ATA region, where large deformations occur, but shows lower performance in the arch and DTA regions, where only relatively small deformations occur.

**Figure 11.**
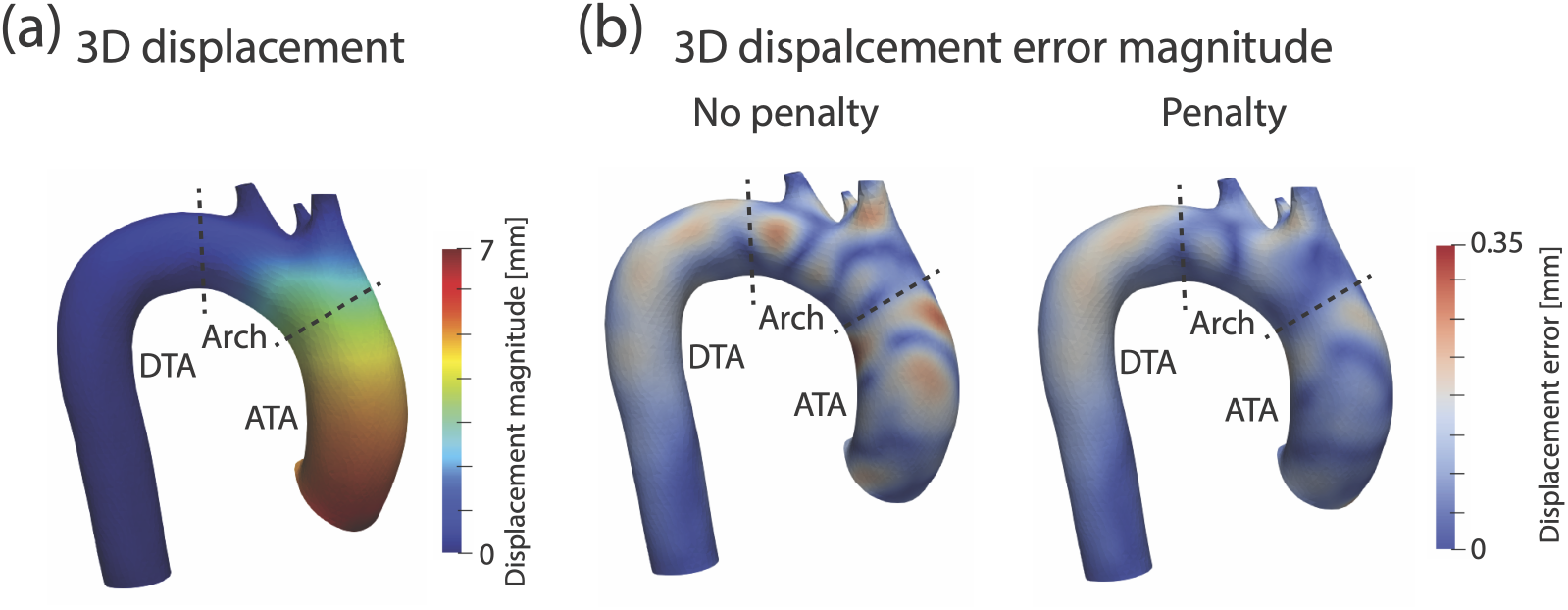
(a) 3D displacement and (b) 3D displacement error with and without penalty parameters for the ATA, arch, and DTA regions.

### 4.6. Limitations

One limitation of this study is the small number of subjects in our cohort. The thresholds used to categorize patients into motion-specific groups were determined using k-means clustering. Because these thresholds (5 and 9 mm) and the corresponding motion-specific penalty parameters were derived from a limited dataset, they may not generalize to larger, real-world cohorts. This study focused on the development of a penalty parameter calibration pipeline. However, it is important to note that these cut-points are consistent with the overall distributions of total displacement reported in a larger study on VDM(D) [12], which included more than 100 patients. Nevertheless, future studies will aim to expand the sample size, allowing for the determination of root motion thresholds and penalty parameters that are more representative of real-world cohorts.

The current VDM(D) can only utilize homogeneous penalty parameters, which means that we can only assign same penalty parameters across the whole aorta. However, our results demonstrate that the values of these parameters correlate with the magnitude of motion. Since the aorta exhibits heterogeneous motion, it would be challenging to find a single set of homogeneous penalty parameters that performs well across all aortic regions.

### 4.7. Future works

Aortic Stiffness Estimation: Once the 3D VDM-derived aortic displacement is available, it is possible to extract multi-axial aortic strain. Assuming a choice of non-linear and multi-axial constitute model, such as four-fiber family model [46], strains can be calculated using a nonlinear rotation-free shell and compared with VDM-derived strains over the entire aorta. This approach would make it possible to determine both axial and circumferential aortic stiffness in a non-invasive way.

## 5. Conclusion

In this study, we successfully calibrated penalty parameters for deformable image registration in VDM(D) using physics-based synthetic data from the rotation-free shell formulation. By tailoring these parameters for each cluster categorized by the magnitude of root motion, we identified that clusters with smaller root motion require larger penalty parameters. These calibrated parameters effectively reduced errors in 3D displacement and multi-axial stretch. Our results demonstrate the importance of customized parameter settings in improving the accuracy of aortic deformation analysis.

